# A model of developmental canalization, applied to human cranial form

**DOI:** 10.1101/2020.10.07.329433

**Authors:** Philipp Mitteroecker, Ekaterina Stansfield

## Abstract

Developmental mechanisms that canalize or compensate perturbations of organismal development (targeted or compensatory growth) are widely considered a prerequisite of individual health and the evolution of complex life, but little is known about the nature of these mechanisms. It is even unclear if and how a “target trajectory” of individual development is encoded in the organism’s genetic-developmental system or, instead, emerges as an epiphenomenon. Here we develop a statistical model of developmental canalization based on an extended autoregressive model. We show that under certain assumptions the strength of canalization and the amount of canalized variance in a population can be estimated, or at least approximated, from longitudinal phenotypic measurements, even if the target trajectories are unobserved. We extend this model to multivariate measures and discuss reifications of the ensuing parameter matrix. We apply these approaches to longitudinal geometric morphometric data on human postnatal craniofacial size and shape as well as to the size of the frontal sinuses. Craniofacial size showed strong developmental canalization during the first 5 years of life, leading to a 50% reduction of cross-sectional size variance, followed by a continual increase in variance during puberty. Frontal sinus size, by contrast, did not show any signs of canalization. Total variance of craniofacial shape decreased slightly until about 5 years of age and increased thereafter. However, different feature of craniofacial shape showed very different developmental dynamics. Whereas the relative dimensions of the nasopharynx showed strong canalization and a reduction of variance throughout postnatal development, facial orientation continually increased in variance. Some of the signals of canalization may owe to independent variation in developmental timing of cranial components, but our results indicate evolved, partly mechanically induced mechanisms of canalization that ensure properly sized upper airways and facial dimensions.

**Author summary:** Developmental mechanisms that canalize or compensate perturbations of organismal development are a prerequisite of individual health and the evolution of complex life. However, surprisingly little is known about these mechanisms, partly because the “target trajectories” of individual development cannot be observed directly. Here we develop a statistical model of developmental canalization that allows one to estimate the strength of canalization and the amount of canalized variance in a population even if the target trajectories are unobserved. We applied these approaches to data on human postnatal craniofacial morphology. Whereas overall craniofacial size was strongly canalized during the first 5 years of age, frontal sinus size did not show any signs of canalization. The relative dimensions of the nasopharynx showed strong canalization and a reduction of variance throughout postnatal development, while other shape features, such as facial orientation, continually increased in variance. Our results indicate evolved, partly mechanically induced mechanisms of canalization that ensure properly sized upper airways and facial dimensions.

## Introduction

The precision and stability with which animal development brings about the adult phenotype has puzzled generations of scientists and is still not well understood. In the 1940s and 50s, the early developmental and theoretical biologist Conrad H. Waddington postulated developmental mechanisms that *canalize* the phenotype in the presence of genetic and environmental perturbations [1–3]. A similar concept of auto-regulatory mechanisms in development was proposed independently and at about the same time by the Russian biologist Ivan Schmalhausen. Today, we know that a disruption of developmental canalization due to environmental or genetic perturbations underlies several of the diseases that are typically considered of multifactorial or unclear aetiology and that can show a spectrum of symptoms [e.g., 4–9]. A related concept in evolutionary biology is “phenotypic robustness,” which refers to the ability of biological systems to produce functional phenotypes in the face of genetic or environmental variation [e.g., 10–12].

In these contexts, the term canalization can refer to phenomena at two different levels, a population pattern and a mechanism in individual development. In population genetics, canalization denotes the reduced effect of an allele, leading to a decrease in genetic variation within a population of individuals [e.g., 12–14]. The second meaning of canalization, to which we refer as “developmental canalization” and which is the focus of this paper, denotes mechanisms that reduce or compensate deviations of *individual development* from its target trajectory. Such deviations can arise from both environmental and genetic perturbations [e.g., 15]. Developmental canalization can reduce phenotypic variation in a population and can (but not necessarily does) underly the population genetic concept of canalization if the canalizing mechanisms are subject to allelic variation.

Development canalization, also referred to as *targeted growth*, has been studied extensively in human and mammalian postnatal growth [e.g., 16–22]. E.g., relatively short individuals tend to undergo accelerated growth, whereas unusually tall individuals show a decreased growth rate to finally achieve an adult body height within the normal range of variation. In human medicine and animal science, these phenomena are also called “catch-up growth” and “catch-down growth,” which are often observed after periods of malnutrition or pathological conditions [e.g., 16, 22, 23]. The term *compensatory growth*, by contrast, usually denotes the developmental compensation of one perturbed trait by another trait, e.g., the hypertrophy of organs or parts of organs after other parts have been removed (some authors, however, use the terms catch-up growth and compensatory growth synonymously). In the same sense, Waddington envisioned development as a “homeorhetic” process: perturbed by environmental or internal fluctuations, individual development varies around a steady trajectory or target trajectory – which he termed the *chreod* (a combination of the Greek words for”determined” or”necessary” and for”pathway”). When the actual trajectory deviates from its target, it tends to return to the chreod due to canalizing processes, as illustrated by his influential metaphor of the *epigenetic landscape* [Fig. 1; see 24, for a review of this concept and it’s impact on modern life science]. Individual development is represented by a rolling marble in this landscape. When it deviates from the valley floor (the chreod or target trajectory), e.g., due to environmental influences, the marble will eventually roll down the wall and return to the floor, until it is pushed away again. The steepness of the wall represents the strength of canalization, the tendency to return to the chreod.

**Figure 1:**
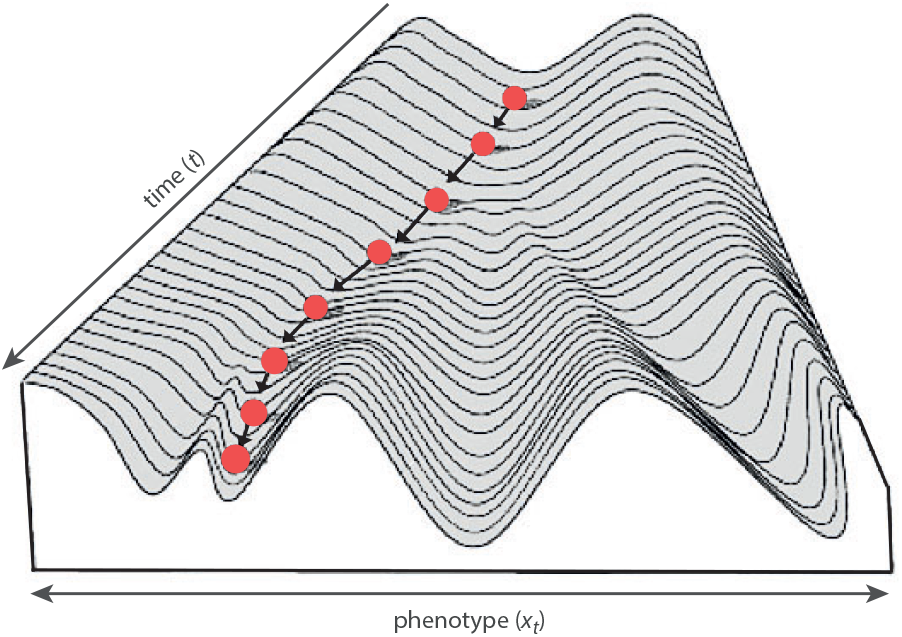
Conrad Waddington’s epigenetic landscape, a landscape over a phenotype space and a time dimension, illustrating the canalization of individual development (the marble) around a target trajectory (the valley floor). Branching valleys represent cell fates, or more generally, predetermined ontogenetic pathways (“chreod”), where the branching points relate to alternative differentiations. The steepness of the walls represent the strength of canalization, the tendency of individual development to return to the chreod. [after 2]

Waddington and many later researchers emphasized the role of canalization in evolution. Canalization can facilitate the accumulation of hidden genetic variation that is “blind” to selection; a breakdown of canalization can release this genetic variation and give rise to episodes of rapid evolution and divergence [e.g., 10–12, 14, 25, 26]. Robustness and canalization may also be subject to evolution themselves [e.g., 27–29]. Although it remains unclear how selection can enhance robustness, the consensus view is that mechanisms canalizing development against environmental and genetic perturbations are a prerequisite for evolvable complex life.

Despite the wide agreement upon the existence of canalizing mechanisms in development, the nature of these mechanisms remains poorly understood. Basically, two lines of explanations and research programs have been proposed [30, 31]. One focuses on the role of particular genes, such as Hsp90 and other chaperones, in buffering the effects of internal and external perturbations of development [9, 32–34]. Quantitative trait loci (QTLs) that affect the variance of a trait but not the trait mean are interpreted as evidence for genetic loci that specifically stabilize the phenotype [e.g., 35, 36]. However, alleles that destabilize development not necessarily indicate an evolved basis of genetic canalization [e.g., 12, 30]. The second line of research explains canalization as an inherent property of the developmental-genetic architecture, such as nonlinear developmental processes, redundancies among pathways, epistatic interactions, averaging effects of independent sources of variation, and epigenetic effects of tissue interactions [30, 37–47].

Most empirical studies on canalization have compared the variance of phenotypic traits across different genotypes or environments. But reduced cross-sectional variance (i.e., variance across individuals of the same age or developmental stage) can result both from increased canalization of developmental perturbations (stronger or faster reversion to the target trajectory) as well as from reduced effects of genetic or environmental variation on development in the first place (i.e., phenotypic development was never perturbed by these factors). Likewise, differences in cross-sectional variance across different age stages not necessarily result from differences in developmental canalization because the variance of newly arising perturbations of development can also change with age. As the target trajectory is usually unobservable and likely to vary across individuals, these phenomena cannot be disentangled by simple comparisons of phenotypic variance. It is even unclear if and how a target trajectory (Waddington’s chreod) is directly encoded in the organism’s genetic-developmental system or, instead, emerges as an epiphenomenon [e.g., 19].

Here, we formalize Waddington’s notion of developmental canalization or targeted growth by an extended autoregressive model. We show that under certain assumptions the strength of canalization and the amount of canalized variance in a population can be estimated, or at least approximated, from longitudinal data (measurements of the same specimens at different age stages), even if the target trajectories are unobserved. We extend this model to multivariate measures and discuss reifications of the ensuing parameter matrix. We apply the univariate and multivariate approaches to longitudinal data on human cranial size and shape.

## Modeling canalized development

Waddington’s view of developmental canalization, as represented by the metaphor of the epigenetic landscape, is closely matched by an Ornstein–Uhlenbeck (OU) process, a mean-reverting continuous-time stochastic process. The Ornstein–Uhlenbeck process, *x*_*t*_, is defined by the following stochastic differential equation:

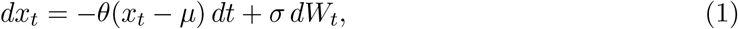

where *θ >* 0 and *σ >* 0. In words, the change of a variable *x* at time *t* is a linear function of the variable’s deviation from *μ*, the *mean reversion level*, plus random noise (a Wiener process). The *mean reversion rate, θ*, quantifies the fraction of the deviation from *μ* that is reverted in the considered time interval. An OU process is a Markov process, that is, the change *dx*_*t*_ depends only on its current value *x*_*t*_, not on any earlier values of *x*.

As a model of individual development, *x*_*t*_ represents a quantitative phenotypic trait at time *t, dx*_*t*_ its developmental change, and *θ* the “strength” by which *x* is attracted by *μ*, which corresponds to the target trajectory or the valley floor of the epigenetic landscape (Waddington’s chreod). *W*_*t*_ represents random and independent perturbations of development. The larger the deviation from the chreod, the faster the reversion (because the reversion is a constant fraction of the perturbation). The slope of the epigenetic landscape thus increases with the distance from the valley floor, giving rise to a concave cross-section of the landscape.

An OU process is a stationary process. In other words, even though *x* fluctuates due to random perturbations, in the long run the mean of *x* approaches the mean reversion level, *μ*, and the variance of *x* stabilizes at 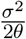. Hence, the variance of the developing trait is determined both by the strength of canalization and the variance of developmental perturbations.

Even though organismal development is a continuous process, it is empirically observed at discrete time points. The discrete-time analog of an OU process is an autoregressive model of order 1, AR(1):

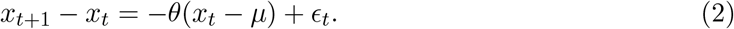

As for the OU process, the parameters *μ* and *θ* correspond to the the target trajectory and the strength of canalization, respectively. Independent random perturbations arising at time *t* are represented by *ϵ*_*t*_. For 0 *< θ <* 2, *x*_*t*+1_ approaches *μ* and reduces the deviation from the target trajectory. Complete canalization occurs for *θ* = 1, whereas 1 *< θ <* 2 corresponds to an overshooting canalization, leading to a damped oscillation of the developmental trajectory around the chreod. For *θ* = 0, the absence of any canalization, developmental perturbations accumulate and *x* shows a random walk.

In the above AR model, the mean reversion level *μ* is constant. For a realistic account of organismal development, however, the target trajectory must change over time and is also likely to vary across individuals unless they are genetically identical (isogenic) and develop in a homogenous environment. Hence, *μ*_*t*_ is not a constant but a variable. Furthermore, empirical studies showed that cross-sectional phenotypic variance can change considerably during mammalian development [e.g., 17, 19, 20, 48, 49]; the strength of canalization may thus differ across developmental stages. In principle, *θ*_*t*_ may also differ among individuals, but if canalization is indeed an inherent property of the genetic-developmental system it is likely to be of similar magnitude in a population. For this reason, and for the sake of a tractable model, we will consider *θ*_*t*_ constant across individuals but free to change over time, giving rise to an extended model:

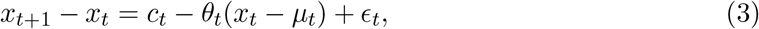

where *c*_*t*_ = *μ*_*t*+1_ *− μ*_*t*_ are the developmental changes of the target trajectories (the “deterministic” component of development). While the developmental perturbations (the “stochastic” component), *ϵ*_*t*_, may be assumed to be uncorrelated across time points (see below for a discussion of this assumption), the developmental changes of the target trajectories can be correlated across time, e.g., when the trajectories continually divergence over a longer period. In this case, the deterministic part can itself be decomposed into a function of its lagged values plus new, independent variation:

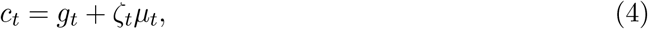

where *g*_*t*_ are deterministic effects on development that newly arise at time *t*, e.g., the effects of developmental genes that are first expressed at this developmental stage. Except for variation in developmental timing (see below), the parameter *ζ*_*t*_ is positive if allelic variation in developmental genes or regulatory factors induces a divergence of growth trajectories. Together, equations 3 and 4 represent a first-order autoregressive moving average process.

In this model, the magnitude of the mean reversion is the same linear function for positive and negative deviations from the chreod, corresponding to a symmetric cross-section of the epigenetic landscape. In sufficiently large samples, the reversion may be estimated as a more complex, non-linear function of the perturbations. Here we will continue modelling the developmental reversion as the fraction *θ*_*t*_ of the perturbation, which can be considered a linear approximation to a nonlinear first-order stochastic process [50].

In the AR(1), canalization is triggered just by the current deviations from the chreod, not by earlier ones. But some developmental perturbations may cause a delayed response, leading to a mean reversion with a longer time lag. This can be represented by an autogressive model of higher order, AR(*p*):

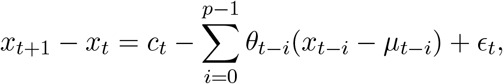

for *p ≤ t*. In words, the current mean reversion is a function *θ*_*t*_ of the deviation at time *t* plus a function *θ*_*t−*1_ of the deviation at time *t −* 1, etc. If not known a priori, the maximal time lag can be estimated from the data, e.g., by the partial autocorrelation function, the partial correlations of *x*_*t*_ with its own lagged values [51, 52].

### Developmental plasticity versus elasticity

The developmental target trajectory may be considered genetically determined, but many allele effects and developmental processes are context-dependent. Non-reversible effects of environmental factors on development, usually referred to as developmental *plasticity*, would then count as a deterministic component of development. By contrast, stochastic short-term perturbations that are subsequently canalized may be considered a developmental *elasticity* [see also 53, section 2.4]. But the same developmental mechanisms that underly the canalization of stochastic short-term perturbations (elasticity) can also contribute the the long-term effect of an environmental factor (plasticity). Similarly, a developmental mechanism that canalizes environmental perturbations may also buffer genetic perturbations of development [e.g., 54–56] so that, again, the same developmental mechanisms can contribute to both the stochastic and deterministic components of development. Hence, even though stochastic and deterministic aspects of development are distinguished in equation 3, this distinction is neither conceptually nor biologically obvious. As shown below, also the separate statistical estimation of these components can be difficult.

### Estimating the strength of canalization

In equation 3, the developmental change, *x*_*t*+1_ *− x*_*t*_, is a linear function, *θ*_*t*_, of the perturbation from the chreod plus newly arising variation: the deterministic changes, *c*_*t*_, and the stochastic perturbations, *ϵ*_*t*_. Write *d*_*t*_ = *x*_*t*_ *− μ*_*t*_ for the developmental perturbations at time *t*, and assume *c*_*t*_, *ϵ*_*t*_, and *d*_*t*_ to be mutually uncorrelated as well as *ϵ*_*t*_ and *d*_*t*_ to be normally distributed with a mean of zero, then the maximum-likelihood estimate of *θ*_*t*_ is given by the least-squares regression of *x*_*t*+1_ *− x*_*t*_ on *d*_*t*_ [e.g., 52]. In the limit of large sample size,

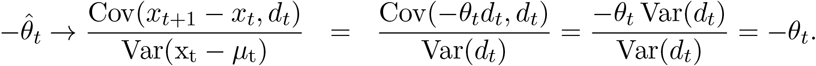

In practice, however, *μ*_*t*_ is unobservable and one can regress the developmental change only on *x*_*t*_ = *μ*_*t*_ + *d*_*t*_ instead of *d*_*t*_. Only if *μ*_*t*_ is constant among individuals (as, perhaps, in an isogenic sample), the ensuing regression slope is an unbiased estimate of *θ*_*t*_. If *μ*_*t*_ varies across individuals, the regression of *x*_*t*+1_ *−x*_*t*_ on *x*_*t*_ yields a biased estimate of *θ*_*t*_, the empirical estimate 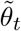:

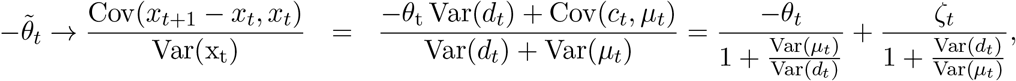

assuming that Cov(*μ*_*t*_, *d*_*t*_) = 0. This shows that the variance of *μ*_*t*_ biases 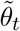 towards zero, whereas long-term trends of the target trajectories (the effect *ζ*_*t*_*μ*_*t*_ on *c*_*t*_) positively bias 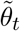. Hence, with increasing (epi)genetic heterogeneity among individuals, the regression of developmental changes on the past trait values increasingly underestimates the strength of canalization.

What is the amount of cross-section variance reduced by canalization? By combining equations 3 and 4, the trait values at time *t* + 1 can be written as

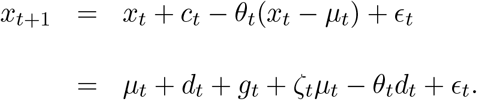

Under the abovementioned independence assumptions, the change in variance from time *t* to *t* + 1 can be decomposed into

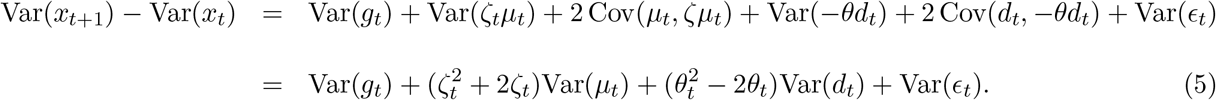

The variance of *x* increases by the variance of new stochastic perturbations, Var(*ϵ*_*t*_), and by the variance of the target trajectories, which can be decomposed in newly occurring “deterministic” variation, Var(*g*_*t*_), and the continuation of earlier divergencies of the target trajectories, 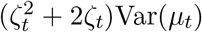. The amount of canalized variance is 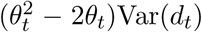. Obviously, the variance reduction is largest for *θ*_*t*_ = 1, when all deviations from the target trajectories are completely reverted (Suppl. Fig. 1). Note that even in the presence of canalization (i.e., 0 *< θ*_*t*_ *<* 2), the cross-sectional variance of *x* can increase if the newly introduced variance exceeds the canalized variance.

Under the same independence assumptions,

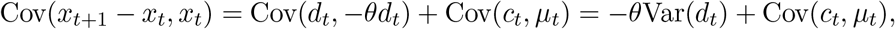

and because 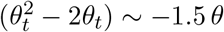 for *−*0.1 *< θ <* 0.6,

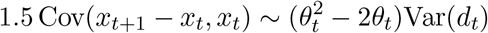

if Cov(*c*_*t*_, *μ*_*t*_) is small (Suppl. Fig. 1). In words, for a small to intermediate canalization coefficient, the canalized variance can be approximated by 1.5 times the covariance between *x*_*t*+1_ and the developmental change *x*_*t*+1_ *− x*_*t*_, as long as newly arising variation among the target trajectories is only weakly associated with the current differences among target trajectories.

In contrast to the estimated strength of canalization, 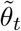, the estimation of the canalized variance is not biased by the variance of *μ*_*t*_. It is also unbiased if the target trajectories all change in the same way (parallel trajectories) or if new differences in target trajectories are unrelated to current differences. However, any long-term trend among target trajectories leads to an underestimation of the canalized variance. For example, if developmental trajectories of body size diverge over a longer time period, large individuals grow more than average compared with smaller individuals. This appears like an “inverse” canalization that amplifies existing individual differences, and 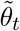 as well as the estimated canalized variance are positively biased.

### Measurement error

In practice, *x* is measured with error. It is well known that measurement error of the independent variable biases the least squares regression slope towards zero [e.g., 57], which also affects the estimation of *θ*_*t*_. But independent error for consecutive measurements of a times series also induces a “regression to the mean” [58, 59]. Write

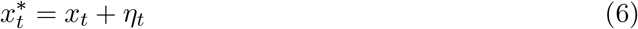

for the measurement of *x*_*t*_ with error *η*_*t*_, and 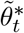 for the empirically estimated *θ*_*t*_ based on 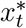. Assuming that measurement error is uncorrelated between time points and with both *x*_*t*_ and *x*_*t*+1_, the estimate equals

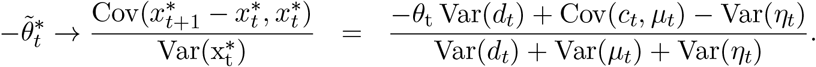

Measurement error decreases the covariance 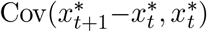 because, intuitively, measurement error at time *t* “disappears” at time *t* + 1. Independent error thus leads to the same statistical signal as canalization. If estimated from repeated measurements, the variance of measurement error can be subtracted from 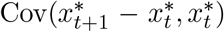 to achieve a more realistic estimate of the canalized variance. Because equations 3, 4, and 6 together constitute a state space model, the parameters can also be estimated by a series of iterations of regression and prediction steps, known as Kalman filter, as long as the target trajectories are relatively stable and close to linear in the observed time interval and the variance of measurement error is known [52, 60, 61].

### Variation in developmental timing

Variation in the onset of rapid growth episodes leads to an increase and subsequent decrease of cross-sectional phenotypic variation (Suppl. Fig. 2). For instance, variation in the onset of the pubertal growth spurt leads to an increased interindividual variance of body height in early adolescence, followed by a decrease of variance after the cessation of growth spurts [e.g. 19]. A very early or late pubertal growth spurt thereby resembles targeted growth. However, such a reduction of variance does not necessarily reflect mechanisms of targeted growth, just variation in developmental timing, which can have a genetic basis or be environmentally triggered. The resulting divergence and subsequent convergence of trajectories can thus be part of the deterministic or the stochastic components of development. In order to avoid this spurious signal of canalization, individual trajectories can be sampled, if possible, at homologous developmental stages rather than at the same ages.

**Figure 2:**
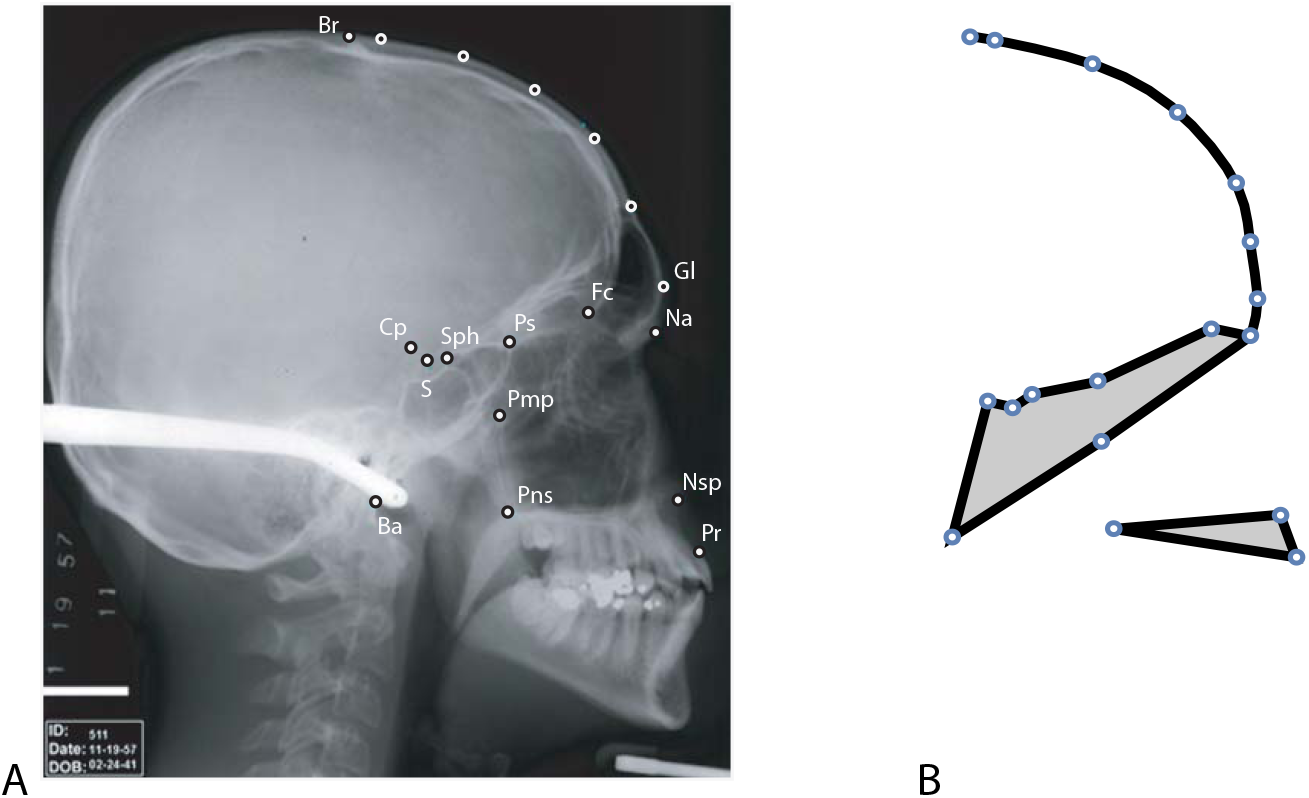
(A) A lateral cranial radiograph of an adult human man with 13 anatomical landmarks: basion (Ba), bregma (Br), clival point (CP), foramen ceacum (FC), nasion (Na), nasospinale (Nsp), greater wings of the sphenoid (Pmp), prosthion (Pr), posterior nasal spine (Pns), planum sphenoideum (PS), sella (S), sphenoidale (Sph), and glabella (Gl). Glabella as well as five points along the frontal bone were treated as semilandmarks; their exact locations were estimated by the sliding landmark algorithm. (B) Mean shape configuration of these 18 landmarks in the sample of 26 individuals, measured from birth to adulthood. The median sagittal outline of the frontal bone is represented by the black curve, and the shape of the cranial base and the maxilla by the two polygons.

### Application to cranial size

We studied developmental canalization in human cranial size using a longitudinal sample of 26 untreated individuals (13 girls, 13 boys) from the Denver Growth Study, a longitudinal X-ray study carried out in the US between 1931 and 1966. On a total of 500 lateral radiographs, covering the age range from birth to early adulthood, 18 landmarks were digitized by E. S. [Fig. 2; for more details see 62]. Six semilandmarks are located on the external outline of the frontal bone, which were allowed to slide along the bone outline so as to minimize the bending energy of the data set around its Procrustes average [63, 64]. Landmark data are available via the DRYAD data repository (XXX). Further information on the Denver Growth Study and the original X-rays are available at https://www.aaoflegacycollection.org/aaof_collection.html?id=UOKDenver. As these are historic and publicly available data, no approval of an ethics committee was necessary.

We used the centroid size of these landmarks (square root of the summed squared distances between the landmarks and their centroid) as a measure of cranial size. Additionally, the area of the lateral projection of the frontal sinus (hereafter, frontal sinus size) was measured on every radiograph. Because the radiographs were not taken at exactly the same age for all individuals, we estimated individual cranial size and frontal sinus size at yearly intervals (2–17 years of age) by unweighted local linear regressions within moving age windows of 2.5 years for each individual. Changing the width of the age window or using a moving average algorithm instead of local linear regression affected the smoothness of the trajectories but lead to qualitatively similar results.

We estimated measurement error by measuring eight age stages for four individuals four times each. Because centroid size is a composite variable of all the landmark coordinates, its measurement error tends to be low. The variance of repeated measurements was in the order of 1-2% of between-individual variation for the different ages. Pooled over all age stages, the variance of repeated measurements was 81.8 (compare to the scale in Fig. 4). The magnitude of measurement error did not correlate with age.

**Figure 3:**
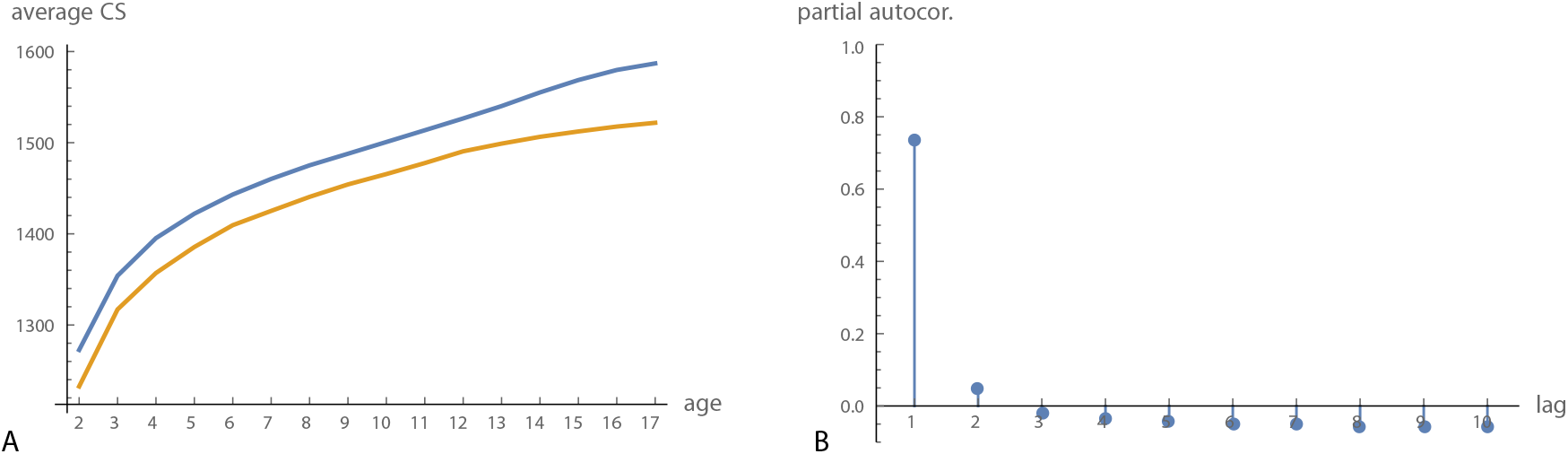
(A) Average cranial size (centroid size) for boys (blue) and girls (orange) in the Denver growth sample. (B) Partial autocorrelation function for cranial size.

**Figure 4:**
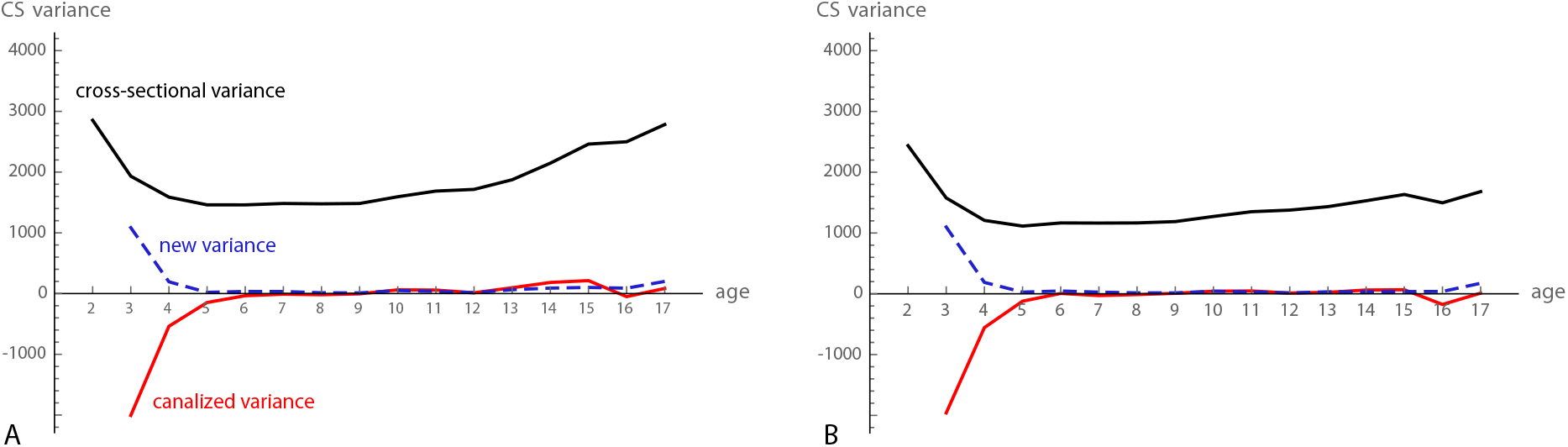
(A) Cross-sectional variance (black curve), canalized variance (red curve), and newly added varaince for cranial centroid size (CS). (B) Same as in A, but here average sexual dimorphism was removed from the data prior to the analysis.

Average cranial size increased fastest until approximately five years of age. Sexual dimorphism in cranial size was relatively constant from 2 to 12 years of age and increased with the beginning of puberty (Fig. 3A). The partial autocorrelation function indicated that only a time lag of 1 is relevant (Fig. 3B), i.e., if conditioning on *x*_*t*_, the earlier observations, *x*_*t−*1_, *x*_*t−*2_, …, did not show a relevant correlation with *x*_*t*+1_. We thus applied an autoregressive model of order 1 to these data. Iterative ARMA estimation and Kalman filter did not perform well, presumably because of relatively unstable, nonlinear growth trajectories.

The cross-sectional variance in cranial size was approximately halved from 2 to 5 years of age and started to increase again at about 10 years. The canalized variance – estimated as 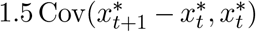– was high in the first few years and decreased to about zero at six years of age (Fig. 4A). From 13 to 16 years, the “canalized” variance was slightly positive, reflecting the continual divergence of male and female growth trajectories. If removing sexual dimorphism from the data by subtracting the age- and sex-specific average from all individuals, this effect goes away (Fig. 4B). The weak signal of canalization at age 16 coincides with the pubertal growth spurt. Because measurement error was negligible for these data, we did not correct for it.

The estimated strength of canalization 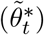 was in the order of 0.5 at age 2–3 and approached zero at about 6 years (Fig. 5). The estimates 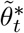 are biased by interindividual variation in the target trajectories. However, correcting for sexual dimorphism, which removes a considerable fraction of variance in target trajectories, only weakly affected the estimated strength of canalization (solid versus dashed lines in Fig. 5).

**Figure 5:**
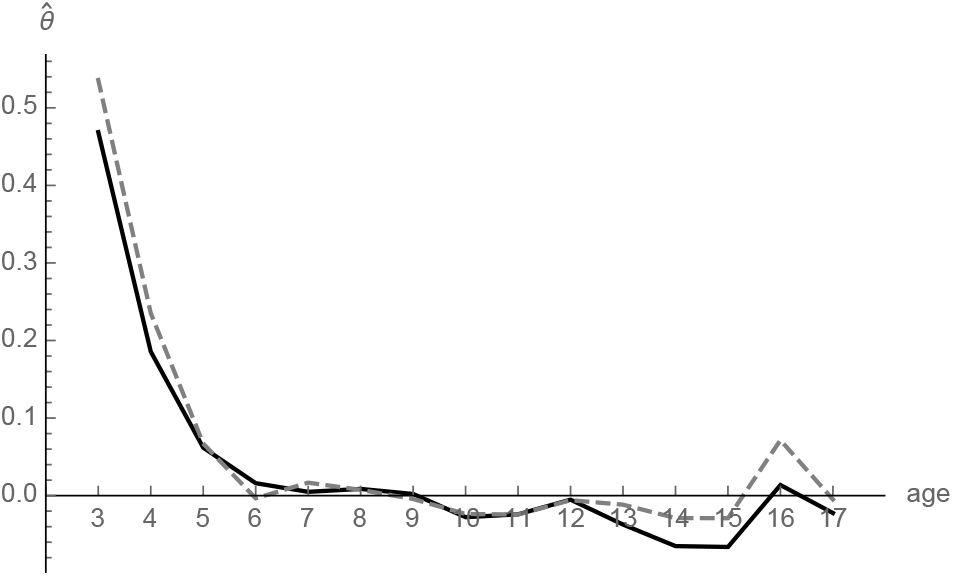
The estimated strength of canalization 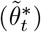 for cranial size before and after removing sexual dimorphism (solid and dashed lines, respectively).

We performed the same analyses also for frontal sinus size. Average fontal sinus size as well as the variance in size increased throughout the studied age range (Fig. 6). The partial autocorrelation function indicated that only a time lag of 1 is relevant. In contrast to cranial size, frontal sinus size did not show any signs of developmental canalization. The estimated canalized variance was even slightly positive, indicating a positive covariance between the increase of frontal sinus size and previous size. Hence, individuals seem to have a tendency to develop small or large frontal sinuses early on. Removing sexual dimorphism slightly lowered all three variance components (not shown).

**Figure 6:**
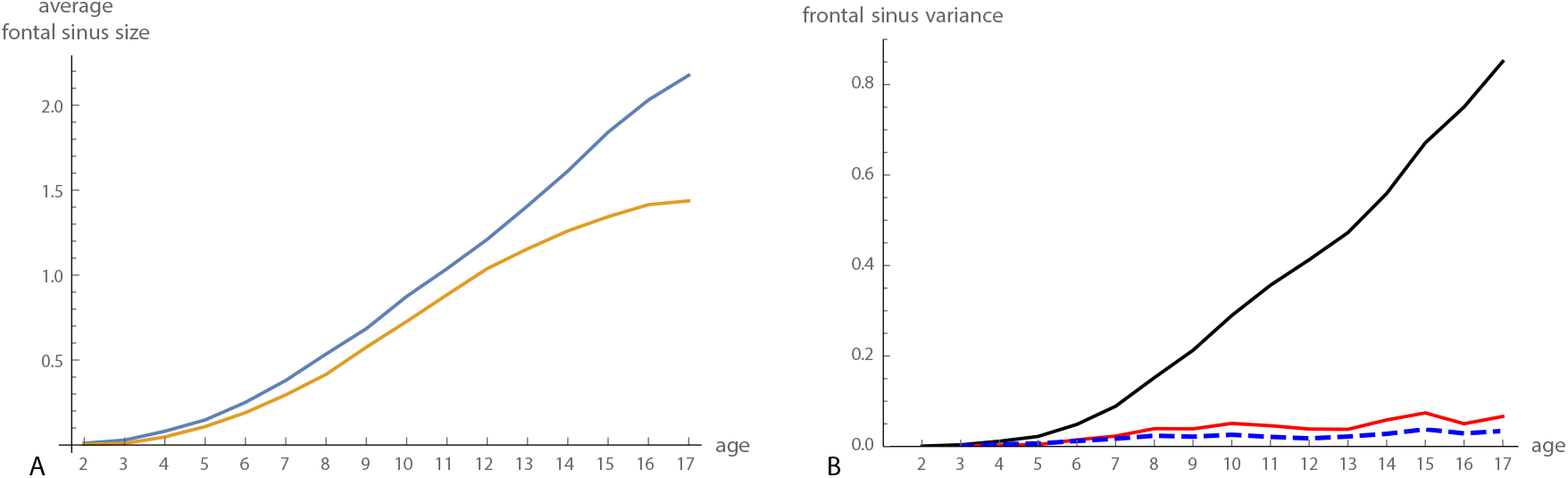
(A) Average frontal sinus size for boys (blue) and girls (orange) in the Denver growth sample. (B) Cross-sectional variance (black curve), canalized variance (red curve), and newly added variance (blue dashed curve) for frontal sinus size.

## A multivariate model of canalization

The univariate model in equation 3 can be extended to multivariate traits:

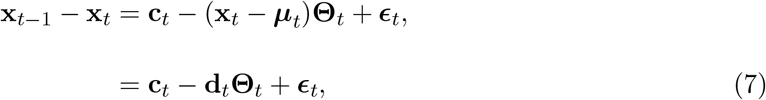

where **x**_*t*_, ***μ****t*, **c**_*t*_, **d**_*t*_, and ϵ_*t*_ are *p*-dimensional random vectors of trait values, chreod values, changes of the chreod values, developmental perturbations from the chreods, and newly arising perturbations at time *t*, respectively. Note that the effect of **x**_*t*_ on developmental change is represented by the *p × p* matrix **Θ**_*t*_, instead of a scalar as in the univariate case. The diagonal elements, Θ_*t,ii*_, quantify how the *i*-th variable at time *t* affects its developmental change from *t* to *t* + 1, conditioned on the other *p −* 1 variables. The off-diagonal elements, Θ_*t,ij*_, quantify how the *i*-the variable at time *t* affects the developmental change of the *j*-th variable. In this sense, they reflect the strength by which one variables *compensates* the perturbation of another variable. These elements don’t have a correspondence in Waddington’s epigenetic landscape.

In the absence of measurement error and variation between target trajectories and if ϵ_*t*_ is multivariate normal with the zero vector as mean, the multivariate multiple regression of **x**_*t−*1_ *−* **x**_*t*_ on **x**_*t*_ yields an unbiased estimate of **Θ**_*t*_. For notational convenience, let **x**_*t*_ and **x**_*t*+1_ be mean-centered, then in the limit of large sample size

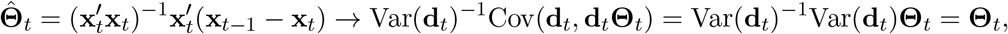

assuming that **c**_*t*_, **d**_*t*_, and ϵ_*t*+1_ are mutually uncorrelated (which is increasingly unrealistic as *p* increases) and that Var(**d**_*t*_) is invertible. As for the univariate case, variation between target trajectories and measurement error bias the empirical estimate 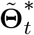, and the regression of 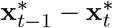 on 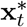 generally underestimates the strength of canalization.

Extending equation 5 to the multivariate model shows that the developmental change in variance-covariance structure due to canalization is

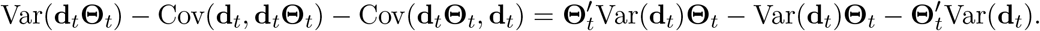

A common multivariate measure of total variance is the sum of the variances of all variables, Tr(Var(**x**_*t*_)). The developmental change in total cross-sectional variance due to canalization is

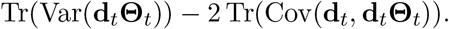

If the elements of **Θ**_*t*_ are small, the quadratic form 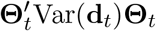 Var(**d**_*t*_)**Θ**_*t*_ is negligible, and the developmental change in variance-covariance structure is mainly determined by Var(**d**_*t*_)**Θ**_*t*_ and its transpose. Because

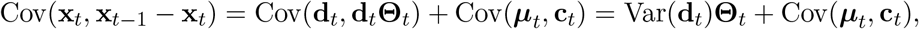

the effects of canalization and compensation on the cross-sectional variance-covariance structure can be approximated by the cross-covariance matrix Cov(**x**_*t*_, **x**_*t−*1_ *−* **x**_*t*_) if the strengths of canalization and compensation are small and the changes of the target trajectories only weakly related to their current states, and if the compensatory effects are approximately symmetric (i.e., Θ_*t,ij*_ *≈* Θ_*t,ji*_).

In many morphometric contexts (e.g., for landmark shape coordinates, collections of inter-landmark distances, or voxel gray values), the measured variables cannot be separately biologically interpreted, and biometric analyses are usually based on linear combinations of the variables. For such data, the elements of **Θ**_*t*_ bear no biological meaning by themselves and are typically too numerous for inspection. Hence, **Θ**_*t*_ may be expressed in its canonical form:

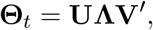

where **U** and **V** are *p×p* matrices of left and right singular vectors, and **Λ** is a diagonal matrix of singular values. The singular vectors can be interpreted as multivariate traits (axes in phenotype space) with maximal canalization (regression slopes). For instance, the regression of the linear combination (**x**_*t*+1_ *−* **x**_*t*_)**v**_1_ on **x**_*t*_**u**_1_ has the slope Λ_1,1_, where **u**_1_ and **v**_1_ is the first pair of singular vectors of **Θ**_*t*_ and Λ_1,1_ the largest singular value. This decomposition of **Θ**_*t*_ allows one to reduce the measured variables to a small set of linear combinations, which can be interpreted as latent variables underlying the observed patterns of canalization and compensation. In the multivariate statistical literature, this is referred to as reduced rank regression [65, 66].

As the change in variance-covariance structure attributable to canalization is approximately proportional to the cross-covariance matrix between trait values at time *t* and their developmental changes from *t* to *t* + 1, one can also compute linear combinations that maximize canalized variance (instead of the strength of canalization) by the singular value decomposition of the cross-covariance matrix:

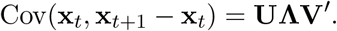

Here, the linear combinations (**x**_*t*+1_ *−* **x**_*t*_)**v**_1_ and **x**_*t*_**u**_1_ have the maximal covariance Λ_1,1_ (as a proxy of canalized variance) among all unit vectors **u** and **v**. The second pair of linear combinations, (**x**_*t*+1_ *−* **x**_*t*_)**v**_2_ and **x**_*t*_**u**_2_, is orthogonal to the first pair and has the second highest covariance, and similarly for subsequent pairs. This approach is equivalent to two-block partial least squares analysis (PLS), which is commonly used in morphometrics [e.g., 67–69]. As the singular values are always positive, interpretations in terms of canalization and compensation must be based on the similarity of the singular vectors, **u**_*i*_, **v**_*i*_, as illustrated by the example below. E.g., pure canalization is indicated by 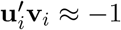, whereas compensation is indicated by 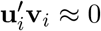.

Note that a stable estimation of 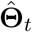 requires a sample size that greatly exceeds the number of variables (*n " p*) because of the matrix inversion of Var(**x**_*t*_). In practical applications, the SVD of the cross-covariance matrix Cov(**x**_*t*_, **x**_*t*+1_ *−* **x**_*t*_) is more reliable, and the associated singular values less affected by sample size than the singular values of 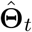.

### Application to craniofacial shape

The 500 configurations of 18 landmarks (Fig. 2) were superimposed by generalized Procrustes analysis in order to standardize for variation in position, orientation, and scale. The resulting 36 shape coordinates were used to quantify craniofacial shape [70–72]. Like for centroid size, we estimated craniofacial shape at yearly intervals by unweighted local linear regressions within a moving age window of 2.5 years for each shape coordinate and each individual separately. Sexual dimorphism was removed for each age group. We computed the cross-covariance matrix between each age group and the subsequent developmental change over one year, 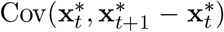, in order to explore the effect of canalization on the cross-sectional variance-covariance pattern. Using different age windows, longer time lags, or the first few principal components of the shape coordinates instead of all 36 coordinates led to very similar results as those presented here.

When estimated by 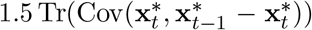, canalization reduced total shape variance until about 6–7 years of age. Cross-sectional shape variance, however, decreased only slightly in this age period, indicating that newly added variance and canalized variance were similar in magnitude (Fig. 7). During puberty, craniofacial shape variance increased rapidly and decreased thereafter, reflecting variation in the timing of pubertal growth episodes.

**Figure 7:**
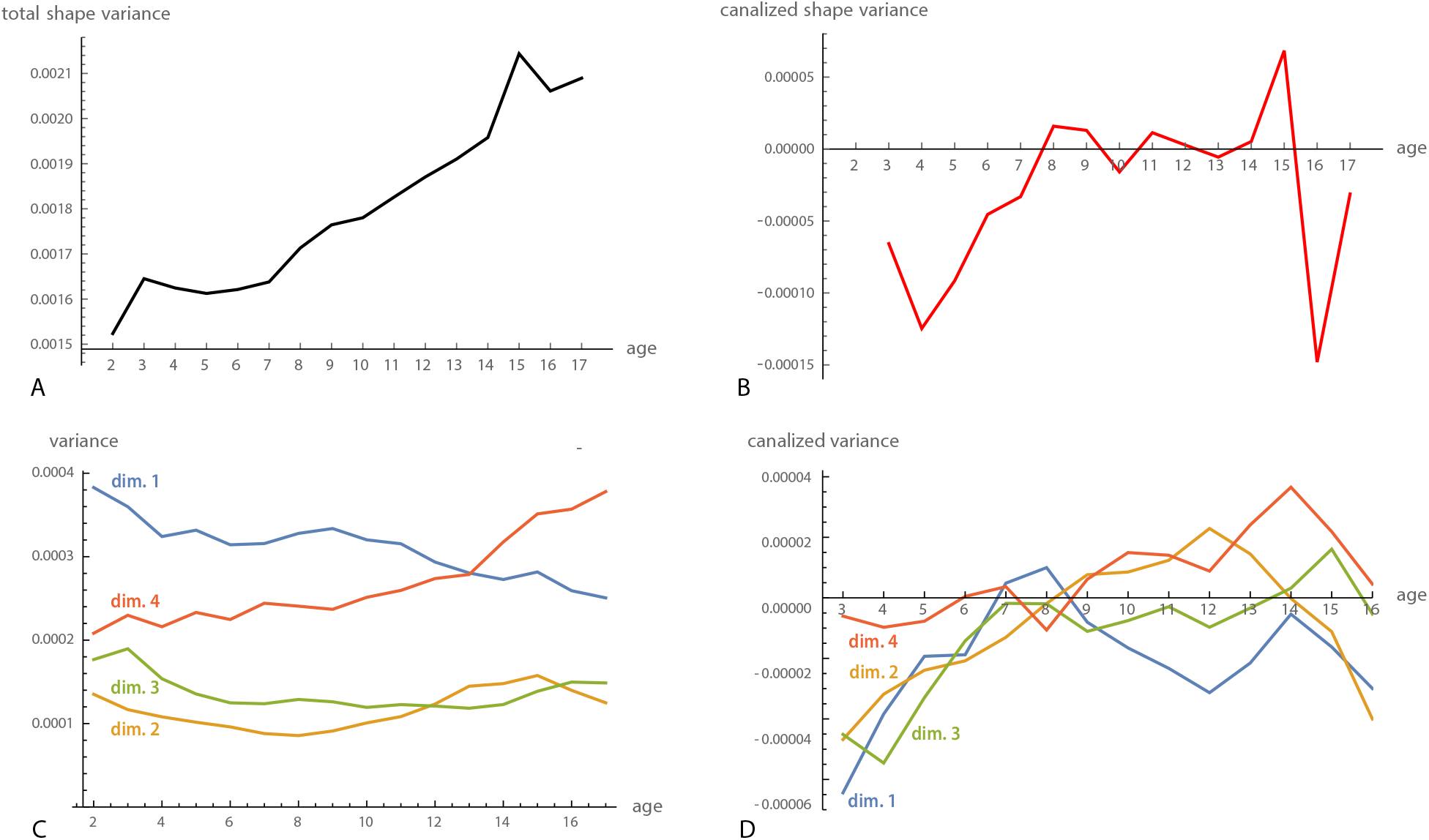
(A) Total craniofacial shape variance from 2 to 17 years of age. (B) Total canalized shape variance, approximated by 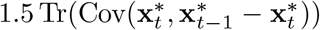. (C) Variance of the first four dimensions of the two-block partial least squares (PLS) analysis between craniofacial shape at age 2 and the shape changes from age 2 to 3 (see Fig. 8). In other words, these are the cross-sectional variances of the shape features with maximal canalization from age 2 to 3. (D) Approximated canalized variance of these four PLS dimensions. The first dimension (blue curve) shows canalization until 6 years and again from 9 to 13, leading to a reduction of cross-sectional variance throughout postnatal development. Dimensions 2 and 3 only show canalization until 6 years of age, whereas dimension 4 shows no signs of canalization (covariances between current shape and subsequent shape change were even positive during puberty, indicating an amplification rather than a reduction of individual differences).

However, pooling the variances of all shape coordinates conceals the heterogeneity of the different aspects on cranial shape [e.g., 49, 73]. Figure 8A shows the first dimension of the PLS analysis between craniofacial shape at age 2 and the shape changes from 2 to 3 years of age (i.e., the first two pairs of singular vectors of 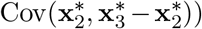, accounting for 50.3% of total squared covariance between craniofacial shape and the subsequent shape change. The first left singular vector, **u**_1_, mainly represented the flexion of the cranial base and the relative size of the maxilla at age 2, which together determine the size of the pharynx. The right singular vector, **v**_1_, was very similar to the left singular vector but in opposite direction, which indicates a pattern of canalization: individuals with a relatively small pharynx at age 2 experience an increased flexion of the cranial base and reduced maxillary growth from age 2 to 3, thus leading to an increase of pharyngeal space. By contrast, individuals with a relatively large pharynx at age 2 show decreased pharyngeal growth. The individual PLS scores, i.e., the linear combinations 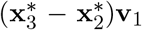 and 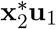, had a correlation of *r* = 0.59 and a regression slope of 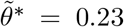. The cross-sectional variances of this shape feature, 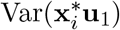, decreased throughout the entire postnatal development (Fig. 7C, D).

**Figure 8:**
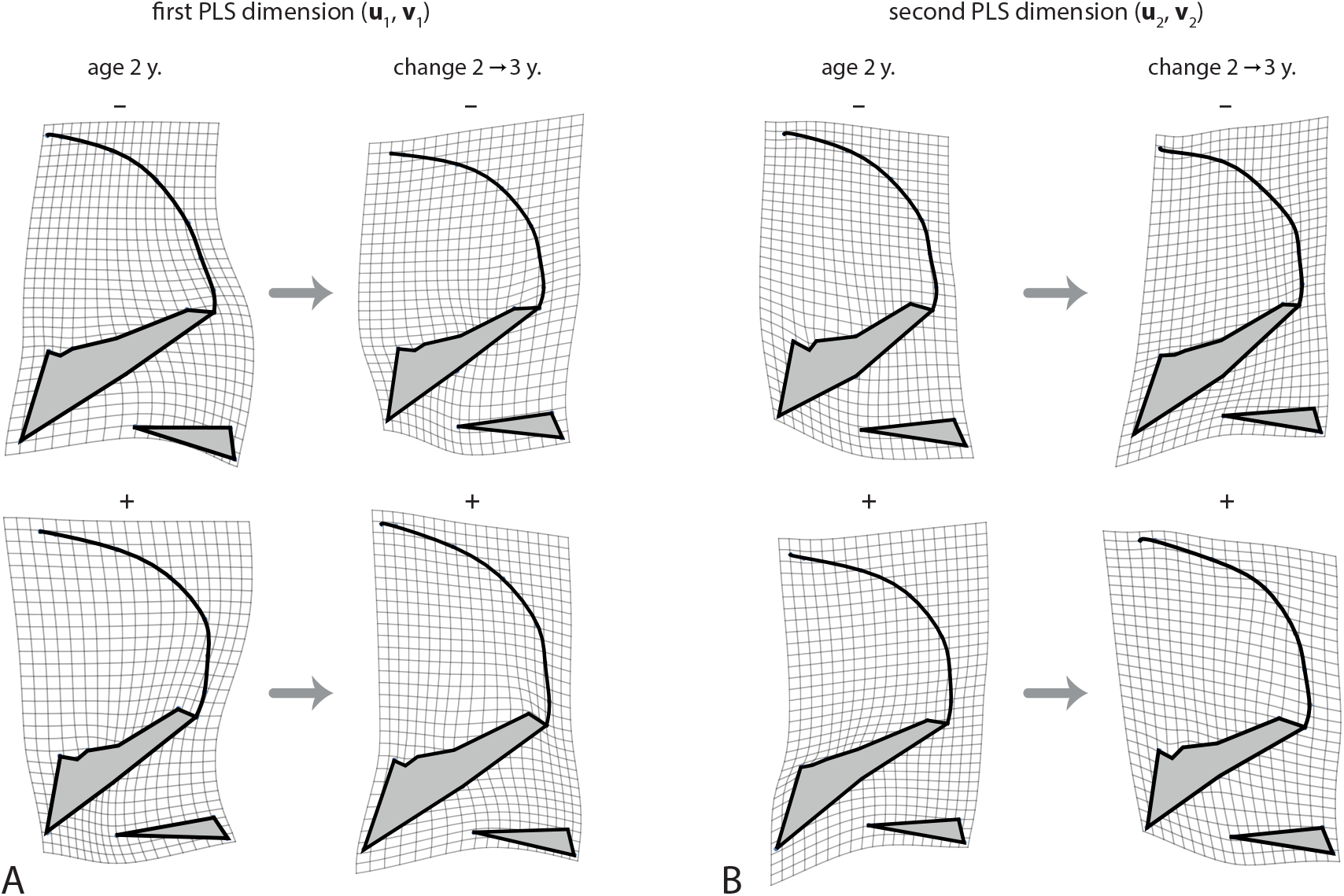
Canalization of craniofacial shape from 2 to 3 years of age was explored by two-block partial least squares (PLS) analysis between the shape variables at age 2 (first block) and the shape changes from age 2 to 3 years (second block). (A) The first PLS dimensions represents the shape pattern with the largest canalized variance. Individuals with a relatively large pharynx due to a retroflexed cranial base and a relatively short maxilla at age 2 (low scores along **u**_1_) tend to experience increased maxillary growth and basicranial flexion from age 2 to 3 years (low scores along **v**_1_). By contrast, individuals with a highly flexed cranial base and relatively long maxilla, both of which constrain the pharynx, tend to undergo a more than average increase in pharyngeal size (high scores along **u**_1_ and **v**_1_). Note that the left deformation grids represent deviations from the mean shape at age 2, whereas the right grids represent deviations from the mean *shape change* form 2 to 3 years. Hence, the maxilla does not shrink in individuals with a large maxilla, but it grows less than in the average growth pattern. (B) The second PLS dimension represents canalization of facial height: individuals with a relatively high face (inferiorly placed maxilla) undergo reduced facial growth, whereas individuals with a relatively low face (including a small pharynx and nasal cavity) show a more than average increase in facial height. Together, this indicates strong canalization of the relative size of the upper airways in early postnatal development.

The second PLS dimension, accounting for 14.9% of total squared covariance, mainly reflected the canalization of relative facial height. Individuals with a relatively high face at age 2 tend to grow less in facial height than individuals with a lower face, and vice versa (Fig. 8B). The correlation between the corresponding scores was 0.70 and the regression slope 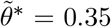. Variance in facial height was canalized during the first 7 years of age and was amplified again during puberty. The third PLS dimension reflected the angulation and relative size of the posterior cranial base, including its effect on relative pharyngeal size (not shown), which was subject to canalization until 6 years of age (Fig. 7C, D). Dimension 4, by contrast, which corresponded to the relative orientation of the face and the pharynx (not its size), did not show any signs of canalization; it even increased considerably in variance during puberty (Fig. 9A, Fig. 7C, D). For the subsequent dimensions, the left and right singular vectors differed in direction and, hence, did not clearly reflect patterns of canalization; also their cross-sectional variances did not decrease (not shown).

**Figure 9:**
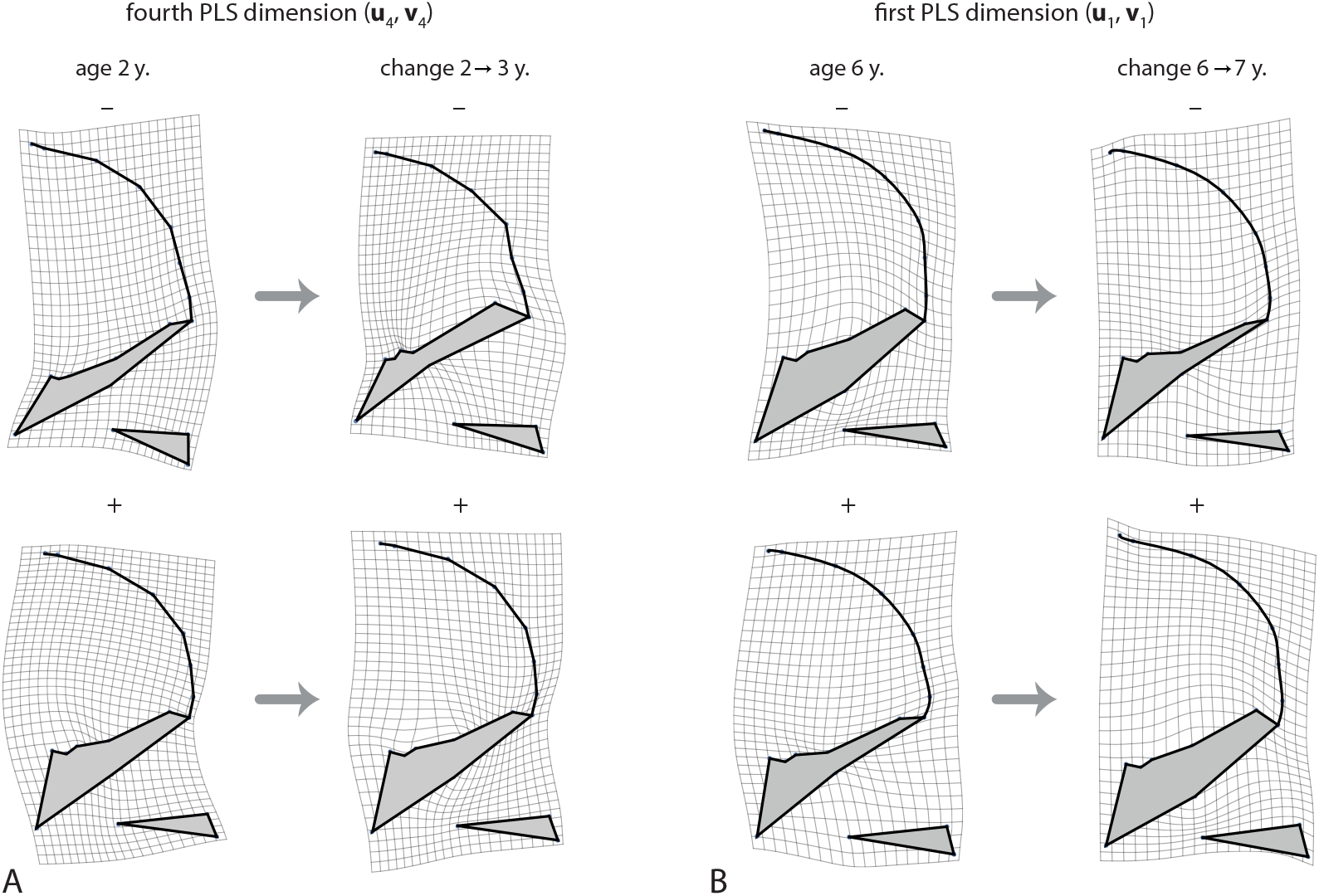
(A) The fourth dimension of the PLS between the shape variables at age 2and the shape changes from age 2 to 3 years. This dimension reflects the relative orientation of the face and posterior cranial base, but unlike the previous dimensions these shape features are not canalized; existing individual differences are even amplified. (B) Canalization of craniofacial shape from 6 to 7 years of age (first PLS dimension). Individuals with a relatively thick cranial base and, therefore, constrained upper pharynx at age 6 undergo reduced thickening of the cranial base, whereas individuals with a thin cranial based show increased growth in this area. Again, this indicates canalization of the relative dimensions of the upper airways.

From about 5 to 10 years, the corresponding PLS analyses revealed another pattern of canalization, mainly affecting the thickness of the cranial base and, indirectly, also the dimensions of the upper pharynx (Fig 9B). Several craniofacial shape features showed strong canalization from 15 to 16 years of age, which likely owes to variation in the timing of pubertal growth periods (see also Fig. 7).

## Discussion

Developmental canalization, or targeted growth, is a prominent but incompletely understood phenomenon in auxology, developmental and evolutionary biology. Most biological studies compared the cross-sectional variance of phenotypic traits across age stages, genotypes, or environmental conditions to infer conclusions about stabilizing processes during development or buffering effects of gene products. In medicine and animal science, by contrast, targeted growth has been described as catch-up growth after periods of retardation in individual growth curves. Both approaches, however, only allow for indirect inferences about developmental canalization because the target trajectory, or Waddington’s chreod, is usually unobservable. It is even unclear if such a “target” of individual growth is explicitly encoded in the organism or emerges as an epiphenomenon. If individual growth is observed at a high temporal resolution over a relatively stable period of development, the individual target trajectories and the perturbations from it may be estimable in a single step by a Kalman filter or related techniques. However, for many organisms and traits, including our data, this is difficult.

We showed that, under certain assumptions, the strength of developmental canalization as well as the canalized variance can still be approximated by the regression slope and the covariance, respectively, between a trait value and its subsequent developmental change in a longitudinal sample. The strength of canalization corresponds to the slope in Waddington’s epigenetic landscape and reflects the ability or propensity of perturbed individual growth to return to the individual’s target trajectory. The canalized variance, by contrast, is the actual average squared reduction of developmental perturbations in the observed sample. Hence, a strongly canalized trait can show little canalized variance if its growth is widely unperturbed.

These estimations rest on the assumption that the developmental perturbations (*d*_*t*_, *ϵ*_*t*_) are statistically independent of the target trajectories (*μ*_*t*_, *c*_*t*_). This assumption seems plausible unless individuals with particular trait values or alleles are considerably more susceptible to environmental or genetic perturbations than other individuals. A second crucial assumption is that the developmental perturbations are uncorrelated across time points. For observations close in time this assumption does not necessarily hold because environmental influences or the expression of particular genes may persist. Positive correlations between perturbations would lead to an underestimation of canalization, but this bias can be reduced by keeping the time lag between observations sufficiently long.

The estimated strength of canalization is biased by inter-individual variation across target trajectories (eq. 5). For isogenic samples, e.g., inbred strains of lab mice, this bias may be minimal because the individuals can be assumed to have very similar target trajectories. In natural populations, including humans, the strength of canalization is underestimated. In our analysis we found that correcting for sexual dimorphism, which considerable reduces variation among target trajectories, only weakly affected the estimates of canalization. Hence, strong signals of canalization are likely to be identifiable also in natural populations.

The estimation of the canalized variance is more robust than that of the strength of canalization. It is not biased by variation among target trajectories, only by the covariance between newly arising differences among target trajectories and their previous states (e.g., long-term divergences of trajectories). In isogenic samples this covariance is close to zero, and in natural populations it decays with the length of the time lag between observations.

In summary, even if the target trajectories cannot be observed or modelled, the strength of canalization can nonetheless be estimated if the genetic and environmental heterogeneity between individuals is small. The canalized variance can even be estimated in natural populations if the time lags between the observations are sufficiently long. As measurement error biases both estimates, it is important to keep the error small and constant across time points and groups, or to estimate it by repeated measurements.

The study of developmental canalization for multivariate traits goes beyond the epigenetic landscape metaphor. For a single variable the strength of canalization is represented by the parameter *θ*_*t*_, whereas its representation for *p* variables requires a *p× p* matrix **Θ**_*t*_, comprising *p* canalization parameters and *p*(*p −* 1) *compensation* parameters. For a small number of biologically meaningful measurements, the estimates of these parameters can be directly interpreted. But for larger numbers of variables, or for measurements that cannot be biologically interpreted (shape coordinates, voxel gray values), the matrix **Θ**_*t*_ must be decomposed into interpretable linear combinations. Multivariate studies of canalization, therefore, may mostly focus on the exploratory identification of features with strong or weak canalization rather than on a precise estimate of the strength of canalization. Linear combinations with maximal strength of canalization can be computed by a reduced-rank regression between the trait values and their subsequent change. Computationally more stable, however, is partial least squares analysis, which yields linear combinations with maximal *covariance* between the trait values and their subsequent change, which is approximately proportional to the canalized variance. Isotropic measurement error biases the covariances but has little effect on the extracted linear combinations.

### Developmental canalization of human craniofacial form

We found that craniofacial size is strongly canalized during the first five years of postnatal development. Canalization decays thereafter and shows a small peak again during puberty, which presumably owes to variation in the timing of pubertal growth spurts. As a result, cross-sectional variance in craniofacial size almost halves between 2 and 5 years of age and continually increases thereafter. Decreasing phenotypic variance of cranial size and body size during postnatal development as well as catch-up growth after periods of retardation have also been reported for many other vertebrates and even insects [e.g., 19, 20, 22, 74–76]. Quantitative genetic studies found that most of the reduced variance is attributable to environmental and maternal effects, whereas additive genetic variance reduces less or even increases during postnatal development [17, 20, 48, 77–79], suggesting that variation in target trajectories has a strong genetic component.

We contrasted the developmental dynamics of craniofacial size with that of frontal sinus size (as inferred from the area of the lateral X-ray projections), which is known to be highly variable in adults [e.g., 80, 81]. For the oldest age group in our data, the coefficient of variation (standard deviation/mean) was 0.51 for frontal sinus size but only 0.03 for craniofacial size. The variance of frontal sinus size increased continually in the observed age range, but the coefficient of variation decreased from 4.0 at 2 years of age to 1.0 at age 6. However, this rapid decrease of the coefficient of variation does *not* imply developmental canalization: we found no signs of canalization for frontal sinus size. The estimated strengths of canalization were even slightly negative, indicating that early individual differences were amplified (rather than canalized) during postnatal development. Presumably, the sinuses lack any canalization because they “grow” only by osteoclast activity and their exact size is of little functional relevance [82]. The coefficient of variation for sinus size just decreased because the mean size increased faster during development than the standard deviation [cf. 47, 83]. Several earlier studies on targeted growth relied on the comparison of coefficients of variation across age groups, but these results should be interpreted with care.

Total variance of craniofacial shape (summed variances of all shape coordinates) slightly decreased from 3 to 5 years of age and then increased by about 30% until age 17. Postnatal reduction of cranial shape variance is not specific to humans; it was also found in other mammals [19, 84–87]. However, such an overall assessment of shape variance partly depends on the multivariate measure of variance and conceals the fact that different shape features (linear combinations of shape variables) can deviate considerably in their variational dynamics [e.g., 49, 73, 88]. We used PLS analyses between the shape variables and their subsequent developmental change over one year to identify canalized features of craniofacial shape. We found multiple aspects of cranial shape that show strong canalization during postnatal development, particularly the flexion and thickness of the cranial base as well as the relative size and position of the upper jaw, all of which determine the dimensions of the nasopharynx. Other features, such as the orientation of the entire face and posterior cranial base, showed no canalization and continually increased in cross-sectional variance.

In the first few years of human postnatal development, the cranial base flexes and the upper jaw increases in length, both of which constrain the pharynx [e.g., 62, 89, 90]. Accordingly, obstruction of the upper airways is relatively common in early childhood [91]. The precise and coordinated growth of the cranial components forming the pharynx thus is crucial to respiratory function. However, in contrast to the pharynx, most other functional components of the cranium, such as the braincase, the orbits, and the jaws, develop in tight interaction with the encapsulated tissues (brain, eyes, tongue and teeth), which tightly coordinates bone growth [89, 92–94]. This lack of a “functional matrix” in the pharynx may make pharyngeal growth particularly prone to perturbations despite its functional importance. Indeed, we found that relative pharyngeal size (PLS dimension 1) had by far the highest variance at age 2 among all the extracted shape features, but this variance was continually canalized throughout postnatal development (Fig. 7C). By contrast, facial orientation (PLS dimensions 4), which is of little functional importance, continually increased in variance until it became the most variable component of facial shape during puberty and thereafter.

A signal of developmental canalization can result from variation in developmental timing and from episodic growth of the entire structure or of its components [e.g., 19, Suppl. Fig. 2]. But our findings of canalization cannot be completely explained by these phenomena. The canalization of cranial size during early postnatal development was much stronger than that observed during and after puberty, and removing average sexual dimorphism in cranial size had little effect on these signals (Figs. 4, 5). Likewise, the shape features related to relative pharyngeal size showed considerably more canalization in early postnatal life than during puberty. Furthermore, developmental dynamics differed strongly between the different shape features and the age periods of strong canalization did not coincide with the periods to fastest developmental change for the different features. Increasing the time lag for the estimations of *θ*_*t*_ and the canalized variance smoothed the curves but did not lead to qualitatively different results.

In summary, craniofacial size as well as multiple aspects of craniofacial shape show a behaviour that resembles developmental canalization in the sense of an auto-regulated or homeorhetic process. Whether or not this behaviour involves a predefined, genetically encoded target trajectory (Waddinton’s chreod) is still unclear. At least for the cranium, it seems most plausible that the targeted behaviour of development emerges from the epigenetic interaction of the developing tissues together with controlled gene expression. Developing soft tissue, such as the brain, eyes, and tongue, provide a matrix that affects and integrates the growth of the surrounding cranial elements. Likewise, sutural growth mechanically induced by stunted or overshooting growth of the adjacent bones canalizes overall cranial shape despite ample variation in the relative dimensions of the constituting bones [47]. Apparently, the lack of epigenetic growth induction in the upper airways accounts for the hyper-variability of nasopharyngeal dimensions in infancy and early childhood. It remains to be studied which mechanisms (e.g., induction by the air flow) account for the pronounced canalization of the nasopharynx during later postnatal development. Understanding the pattern and timing of developmental canalization in the cranium can also aid the effective timing of orthodontic treatment and the reduction of relapse [e.g., 95, 96]. The methods proposed here will allow for the comparison of developmental canalization and targeted growth across populations, genotypes, treatment and age groups in future research.

## Supporting Information

**Supplementary Figure 1:**
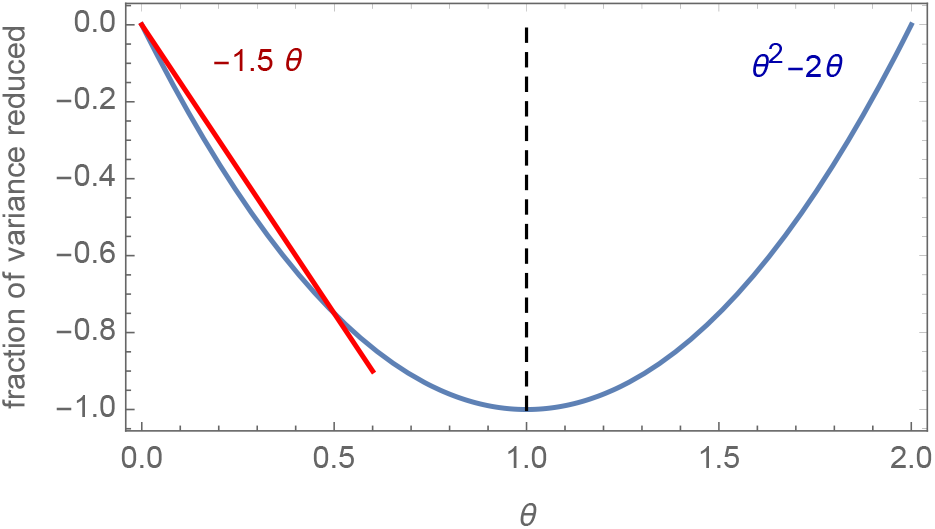
In the autoregressive model of equation 3, the fraction of variance reduced due to canalization equals *θ*^2^ *−* 2*θ*, which is plotted here against *θ*, the strength of canalization. For *θ* = 1 (dashed line), all developmental perturbations are canalized. For 0 *< θ <* 0.6, this function can be approximated by the linear function *−*1.5 *θ* (red line).

**Supplementary Figure 2:**
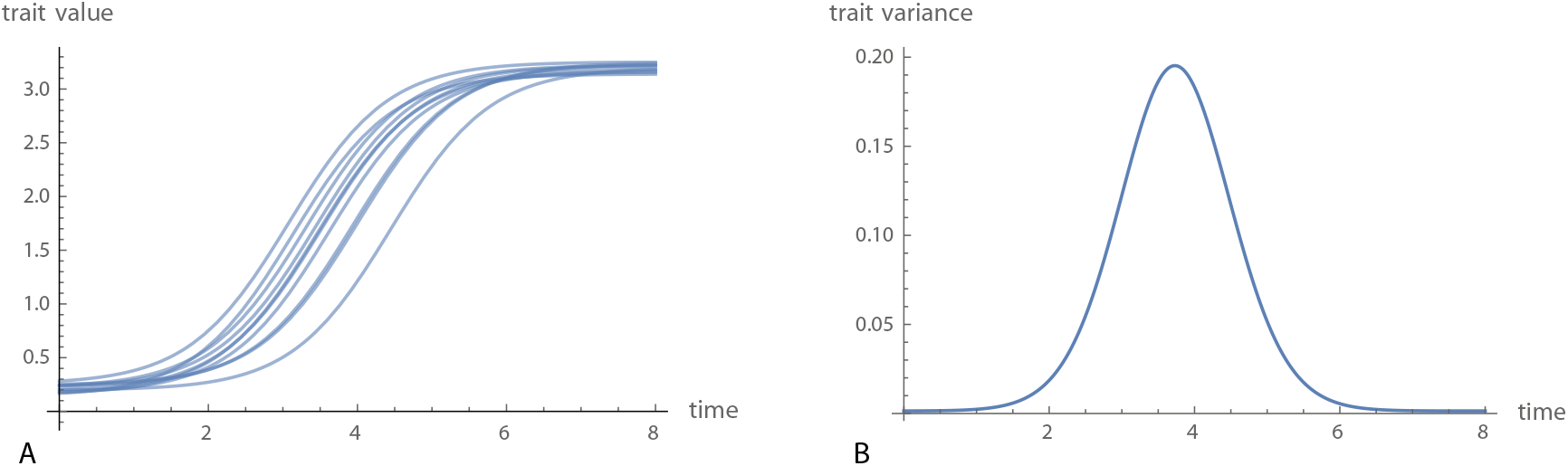
(A) Ten hypothetical growth curves with episodes of rapid growth. The curves have the same shape but differ in the timing of the growth spurt. (B) Due to this variation in timing, the cross-sectional variance of the trait increases and subsequently decreases when all individuals have experienced the growth spurt.

